# Low Cost Metabolic Fuel Sensor for On-Demand Personal Health and Fitness Tracking

**DOI:** 10.1101/2020.02.20.958264

**Authors:** L. M. Candell, R. W. Hoyt, J. M. Mahan, A. M. Siegel, R. M. Standley, K. J. Thompson, G. A. Shaw

## Abstract

Metabolic health in the general population has declined significantly in just one to two generations despite increased emphasis on dieting and exercise. A challenge in prescribing a healthy diet and exercise regimen is the variability individuals exhibit in response to particular foods, calorie restricted diets and exercise regimens. This paper describes a prototype metabolic fuel sensor designed for ease of use and personal tracking of metabolic energy expenditure and fuel substrate utilization. Examples of the sensor measurements and potential applications to weight management and tracking of chronically high blood glucose are described.

## I. Introduction

The cause and effect relationships between diet, metabolism, socioeconomic status, and debilitating medical conditions, such as type-2 diabetes, are a subject of active research and debate [1,2,3,4]. The fact that there is not universal agreement regarding the connections is evident from the multitude of different, often antithetical (e.g., low fat versus low carb), diet plans and the prevalence of degraded or malfunctioning metabolisms among a significant fraction of the US population. According to data published by the CDC in 2016 [5], nearly half of the adults in the US are either pre-diabetic (86 million) or diabetic (29 million) and over 35% of American adults and close to 17% of children, are obese. Recent work by the Weizmann Institute focusing on glycemic control, and employing continuous glucose monitors, has shown that glycemic response to foods can vary significantly across individuals [6]. Their work underscores the importance of glycemic response tracking in order to personalize the nutrition profiles to individual needs and goals.

Just as glycemic response to particular foods varies significantly across individuals, so also the individual response to an exercise-induced hypocaloric diet varies considerably. In a three-month long weight loss experiment involving seven pairs of identical twins [7], a large variance between predicted and actual weight loss across pairs of twins was observed, whereas the weight loss between twins was strongly correlated, presumably due to genetic factors.

The individual variability reported in these and other studies accounts, at least in part, for the lack of success that formulaic weight loss plans and exercise regimens have achieved in reversing the downward trend in metabolic health across the population. Given the individual variability, it is difficult to predict responses to dietary and exercise interventions and thus to formulate an effective regimen for achieving weight loss goals or reversing type-2 diabetes. However, the availability of a low-cost, simple to use sensor for tracking metabolic response to diet and exercise could enable individuals to develop more successful diet and exercise regimens aimed at achieving weight loss and metabolic health goals. Until now, a barrier to personal metabolic tracking has been the absence of a low-cost, easy-to-use sensor capable of tracking metabolic fuel state and energy expenditure over a range of activity.

## II. COBRA Metabolic Fuel Sensor

The Carbon-dioxide Oxygen Breath Respiration Analyzer (COBRA) is a metabolic fuel sensor developed for personal on-demand metabolic measurement. COBRA employs the method of indirect calorimetry to determine energy expenditure (EE) as well as the metabolic fuel mix (relative contributions from carbohydrate oxidation versus fat oxidation). This method relies an accurate measurement of the volume rate of oxygen consumption (VO_2_) and carbon dioxide production (VCO_2_) by an individual. The ratio of VCO_2_ to VO_2_ in exhaled breath, termed the respiratory exchange ratio (RER), is indicative of the average cellular-level respiratory quotient (RQ) and is the key to estimating energy expenditure (EE) and fuel substrate use [8]. In clinical settings, RER is measured by employing a mixing chamber with sufficient volume to collect several exhaled breaths (e.g. 3-4 liters). The contents of the chamber are continuously sampled and passed through gas sensors to measure the residual oxygen and carbon dioxide in the exhaled breath.

Given the COBRA goal of personal use, the over-arching priorities guiding the design of the sensor were small size, weight, and power (SWaP), low cost, ease of use, and low-maintenance. These high-level, qualitative requirements translate into the following measureable design goals:

- Small size, weight and power (SWaP)
- Use of low-cost commercially available components
- Gas sensor lifetimes measured in years
- Stable gas sensor calibration with capability to perform ambient-air calibration checks
- No expendables or consumables required to make a measurement
- Autonomous confirmation of a successful measurement
- Accommodation of the full range of physical activity from resting to VO_2_max measurements
- Wireless operation and smartphone interface
- Battery endurance and on-board memory to support at least one full day of measurements
- Operation over (above freezing) temperature and humidity extremes encountered outdoors

The traditional approach to realizing a mobile metabolic measurement system capable of supporting a range of activity levels has been to eliminate the need for a bulky ~3-4 liter mixing chamber by employing a constant-rate pump to sample a portion of the exhaled breath and to continuously measure the volume fraction of O_2_ and CO_2_ in this side-stream flow. This approach, termed breath-by-breath, requires fast (and hence costly) gas sensors, rapid sampling of the breath profile, and careful calibration of the time delays between the volumetric flow measurement and the sequential measurements of the gas sensors.

To realize the design objectives of both low-cost and small SwaP, the COBRA sensor employs an innovative passive proportional side-stream sampling technique [9][10] that enables the use of slow (low-cost) gas sensors and a mixing chamber volume ~100X smaller than traditional systems. A prototype of the COBRA sensor, employing 3D printed flowtube and polycarbonate machined mixing chamber, is shown in Fig. 1a. The sensor is completely passive with no moving parts other than two inlet umbrella valves. In production, the flowtube and mixing chamber will be injection molded and total weight of the sensor is estimated to be less than 150g. In unit quantities, the parts for the sensor, including the printed circuit board fabrication and population, are under $500. In volume production, the cost of the sensor is anticipated to be in the $200-$300 range, making it affordable for personal use.

**Fig. 1:**
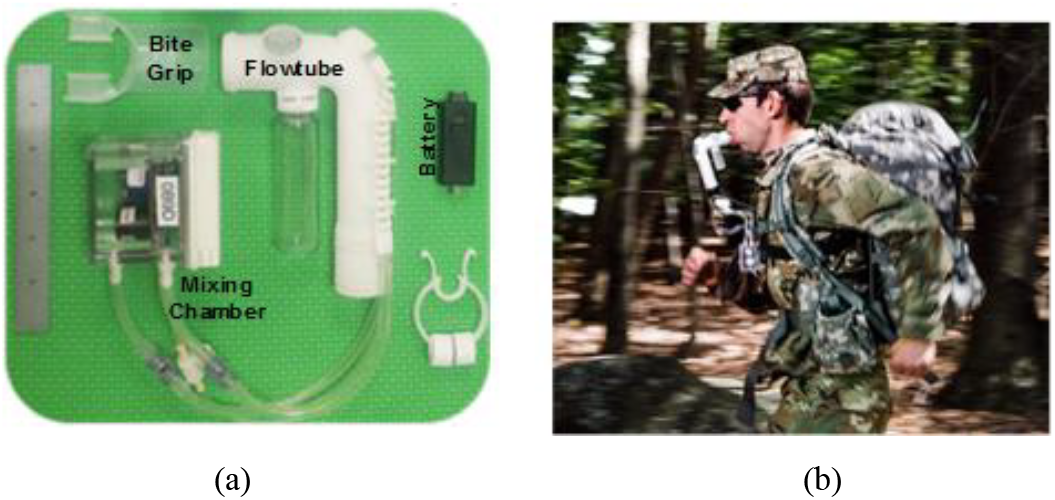
(a) Prototype COBRA sensor (b) hands-free use

The COBRA sensor measures the volume rate of O_2_ consumption and CO_2_ production. Together these parameters enable calculation of the energy expenditure rate of the user as well as the relative proportion of energy derived from fats versus carbohydrates, under the assumption that protein is a minor contributor to metabolic energy. An RER of 0.7 indicates that metabolic energy is being supplied exclusively from fat, whereas an RER =1.0 indicates that metabolic energy is being supplied exclusively from carbohydrates. Values in the range between 0.7 and 1.0 imply a mixture of fat and carbohydrate fuel sources. The RER for protein is nominally 0.85.

## III. Sensor Output And Applications

Fig. 2 shows the energy expenditure, RQ, and volume flow rates measured by the COBRA sensor for three different levels of exercise intensity. Note that low-intensity exercise such as walking is primarily fat burning, while high intensity exercise such as running is primarily carb burning and burns calories at nearly 5x the rate of walking.

**Fig. 2:**
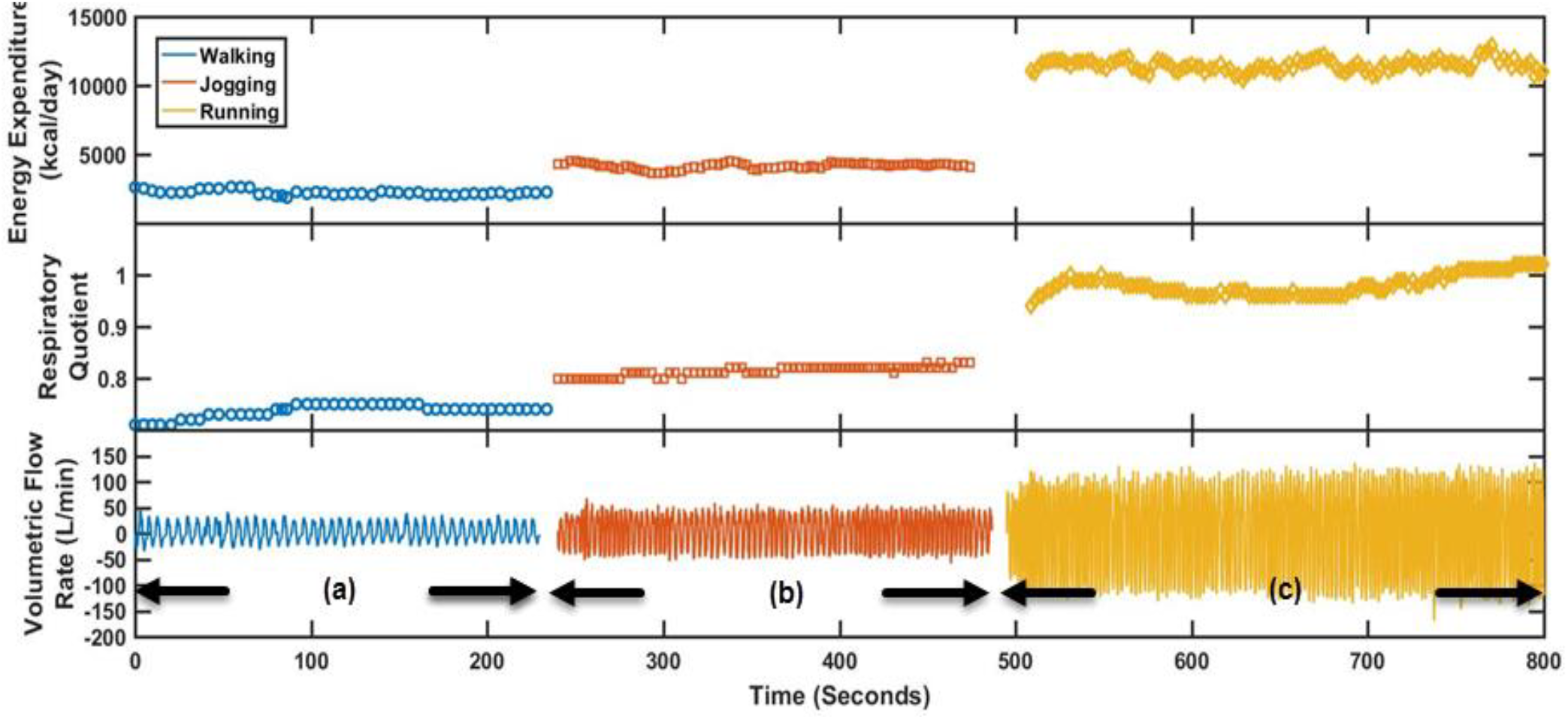
Three different data collects. (a) Walking: RQ ~0.75 & EE~2200kcal/day (b) Jogging: RQ ~0.80 & EE ~4400kcal/day (c) Running: RQ ~ 1.0 & EE ~12000kcal/day.

In the case of walking, the low RQ observed is because walking is dominated by slow-twitch muscular processes which are typically fat burning. In this particular case, the subject has an RQ of 0.75 or a molar mix of approximately 17% carbohydrates and 83% fats. As the subject begins to work harder, more glycogen from the skeletal tissue is used for energy production and the RQ increases to 0.80 and the EE increases from about 91 to 183 kcal/h (2200 to 4400 kcal/day). As the exercise intensity further increases, glucose becomes the dominant fuel and takes over control of the macronutrient selection, raising the RQ to 1. During anaerobic exercise and build up of CO_2_, RER may rise above 1 (lactic acidosis), which is one of the conditions in which RER is not synonymous with cellular RQ.

In any case, these data exemplify the transition in metabolic fuel preference as a function of exercise intensity. The transition points and the values of RQ and EE, paired with the exercise information, can be used to assess individual fitness and track changes in fitness and endurance over time.

### A. Feedback for Tuning Weight Management Protocols

The traditional approach to weight loss is to estimate calories associated with food intake and activity, create an energy deficient diet, and monitor progress toward weight loss goals with a scale. Not surprisingly, this approach fails much of the time for several reasons. For one, counting food calories and estimating activity calories is fraught with error, and secondly, the impact diet and activity choices can be obscured by weight fluctuations of several pounds or more a day unrelated to reduction in adipose tissue.

Rather than a one-size fits all formulaic energy deficit diet for weight loss, the metabolic sensor provides a previously unavailable tool for tracking and tailoring macronutrient intake and exercise activity to increase the likelihood of achieving weight loss and weight management goals. In particular, a on-demand measurement of RQ and EE may prevent dieters from deceiving themselves about the efficacy of daily adjustments in dietary macronutrients or exercise regimen aimed at achieving weigh loss or better glycemic control. The measurement of RQ throughout the day provides quantitative evidence of the impact of dietary and exercise choices on the goal of staying in the fat burning zone (low RQ). In comparison, measured body weight is impacted by numerous confounds unrelated to loss of excess fat, such as hydration and digestive state. Consequently, daily body weight is difficult to correlate with specific diet and exercise choices made during the course of a day. Fig. 3 shows an anecdotal example of how metabolic fuel mix changes in response to diet and exercise. With one exception, the RQ measurements in Figure 3 were all made at rest. For weight loss, diet and exercise choices that result in low resting RQ are preferred since they result in the highest rate of fat burning throughout the day. Note the switch to nearly all carbs in response to the high carb pizza/cookie lunch. The walk at 17:11 is predominantly fat burning and overshadows the nominal 1-2 kcal/min carb burning when at rest. However, while walking burns fat, it does not reduce the excess carbohydrates that prevent fat burning after the walking has ceased. In comparison, even three hours after completing a half marathon on 13 Oct, the depletion of carb stores induced by the endurance run resulted in reliance predominantly on fat burning to meet resting metabolic energy needs.

**Fig. 3:**
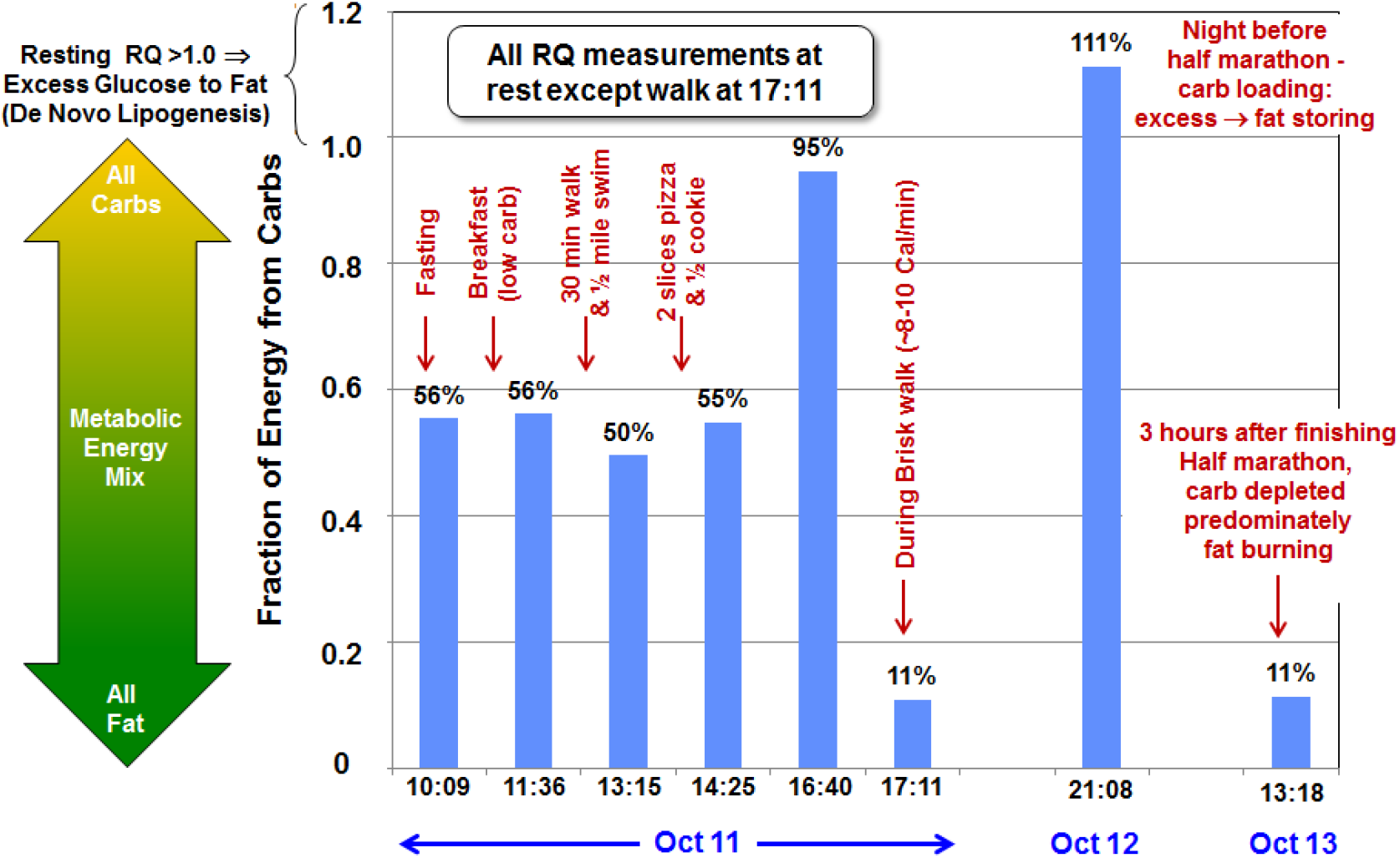
Anecdotal examples of the impact of diet and exercise on fuel substrate mix. Selected measurements from a 3 day period encompassing a high carbohydrate dinner on 12 Oct in preparation for half-marathon race on 13 October.

### B. Type-2 Diabetes and Pre-Diabetes Detection and Intervention

Pre-diabetes is defined by a fasting blood glucose level between 100 – 125 mg/dl. A normally functioning endocrine system strives to keep blood sugar below 100-110 mg/dl at all times and above 80mg/dl to avoid hypoglycemia. A primary mechanism for achieving this regulation in the face of carbohydrate over-consumption is to defer oxidation of fats and preferentially oxidize excess glucose to meet metabolic energy needs. Consequently, a chronically high resting RQ implies a lack of the metabolic flexibility needed to quickly dispose of high glycemic foods. Measurement and tracking of RQ variability may prove to be a non-invasive indicator of the onset of type-2 diabetes. This is one of several intriguing applications of the COBRA sensor that have yet to be investigated in a statistically significant clinical trials.

## IV. Summary And Conclusion

Both young and old members of the general population are exhibiting increasing rates of metabolic syndrome. Extreme diets (e.g., ketogenic) and extreme exercise (e.g. high intensity) have been shown to be effective in achieving weight loss and avoiding or reversing type-2 diabetes, but their extreme nature makes them difficult to sustain for many individuals. An alternative is to provide an individual with the capability to measure energy expenditure and metabolic energy mix on demand, enabling the individual to design a personalized diet and exercise regimen that creates the daily energy deficit, and keeps them in the fat burning zone necessary to achieve weight loss and glycemic control goals.. Until recently, the sensing technology to provide mobile, on-demand metabolic feedback has been limited to resting measurements of energy expenditure, or required sensors of significant size, weight, power and cost.

We have described the development of a prototype low-cost, simple-to-use, personal sensor to provide metabolic awareness to individuals. The sensor runs for a day on a battery charge, archives the data for later fusion with personal sensors, and supports a Bluetooth smartphone interface for control and data display. Initial prototypes of the sensor have undergone an independent validation and found to provide reliable measurement of VO_2_, VCO_2_, energy expenditure and related parameters, over a range of activity levels [11].

Low-rate initial production of the sensor is scheduled for the second quarter of 2019 and we look forward to collaboration with researchers interested in incorporating the sensor into clinical trials and wellness programs.

## ACKNOWLEDGMENT

The authors wish to acknowledge Dr. Karl Friedl for his early recognition of the potential benefits of the sensor for tracking and improving metabolic health and performance, and Ms. Holly McClung and her team at the US Army Research Institute of Environmental Medicine for conducting an independent assessment of COBRA performance and providing valuable feedback on design enhancements.

## Notes

This material is based upon work supported by the Department of the Army, specifically funding from the Defense Health Agency Health Program, under Air Force Contract No. FA8702-15-D-0001. Any opinions, findings, conclusions or recommendations expressed in this material are the private views of the authors and are not to be construed as official or as reflecting the views of the Army or the Department of Defense. Citations of commercial organizations and trade names in this report do not constitute an official Department of the Army endorsement or approval of the products or services of these organizations.

